## Supplementary data for "The M-phase regulatory phosphatase PP2A-B55δ opposes protein kinase A on Arpp19 to initiate meiotic division"

Lemonnier T. *et al.*

**SUPPLEMENTARY DATA**

**Supplementary Figure 1. Cter-GST-Arpp19 is phosphorylated at S109 by PKA and does not affect meiosis resumption.**

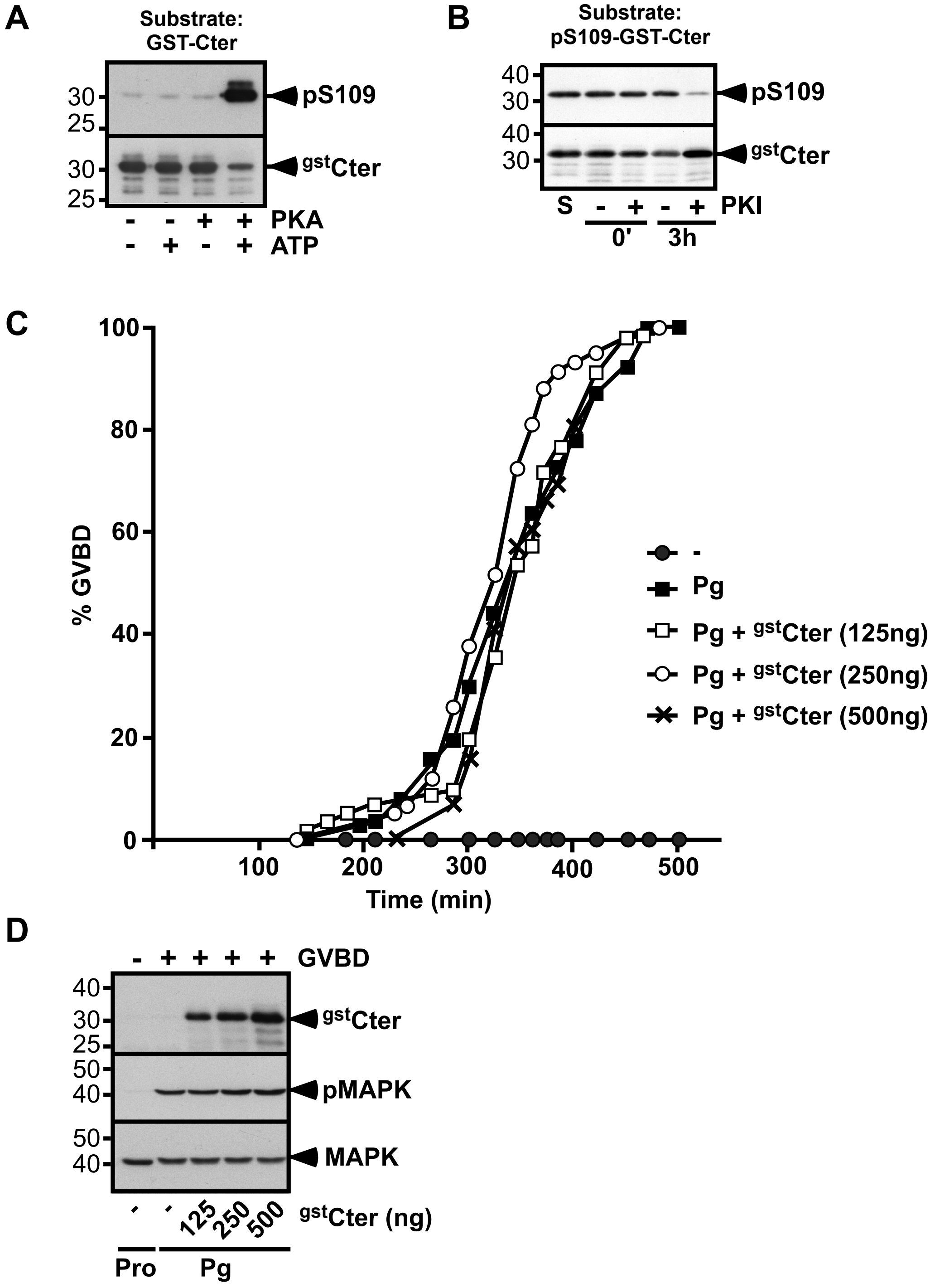

(**A**) Cter-GST-Arpp19 was incubated with or without PKA catalytic subunit in the presence or in the absence of ATP. S109 phosphorylation of Cter-GST-Arpp19 (pS109) and total Cter-GST-Arpp19 ($^{gst}$Cter) were western blotted using respectively phospho-S109-Arpp19 and GST antibodies. (**B**) Cter-GST-Arpp19 was *in vitro* phosphorylated by PKA and then incubated for 3h in prophase extracts, supplemented or not with PKI. S109 phosphorylation of Cter-GST-Arpp19 (pS109) and total Cter-GST-Arpp19 ($^{gst}$Cter) were western blotted using respectively phospho-S109-Arpp19 and GST antibodies. (**C**) Prophase oocytes were injected with various amounts of Cter-GST-Arpp19 ($^{gst}$Cter) as indicated, and stimulated with progesterone (Pg). GVBD was scored as a function of time. (**D**) Same experiment as in (**C**). Prophase (Pro) or progesterone-stimulated oocytes (Pg), injected or not with Cter-GST-Arpp19, were collected at the time of GVBD. Cter-GST-Arpp19 ($^{gst}$Cter) was pulled-down and western blotted with the anti-GST antibody. Total oocyte extracts were western blotted with antibodies against phosphorylated MAPK (pMAPK) and total MAPK.

**Supplementary Figure 2. Protocol of S109-phosphatase biochemical isolation.**

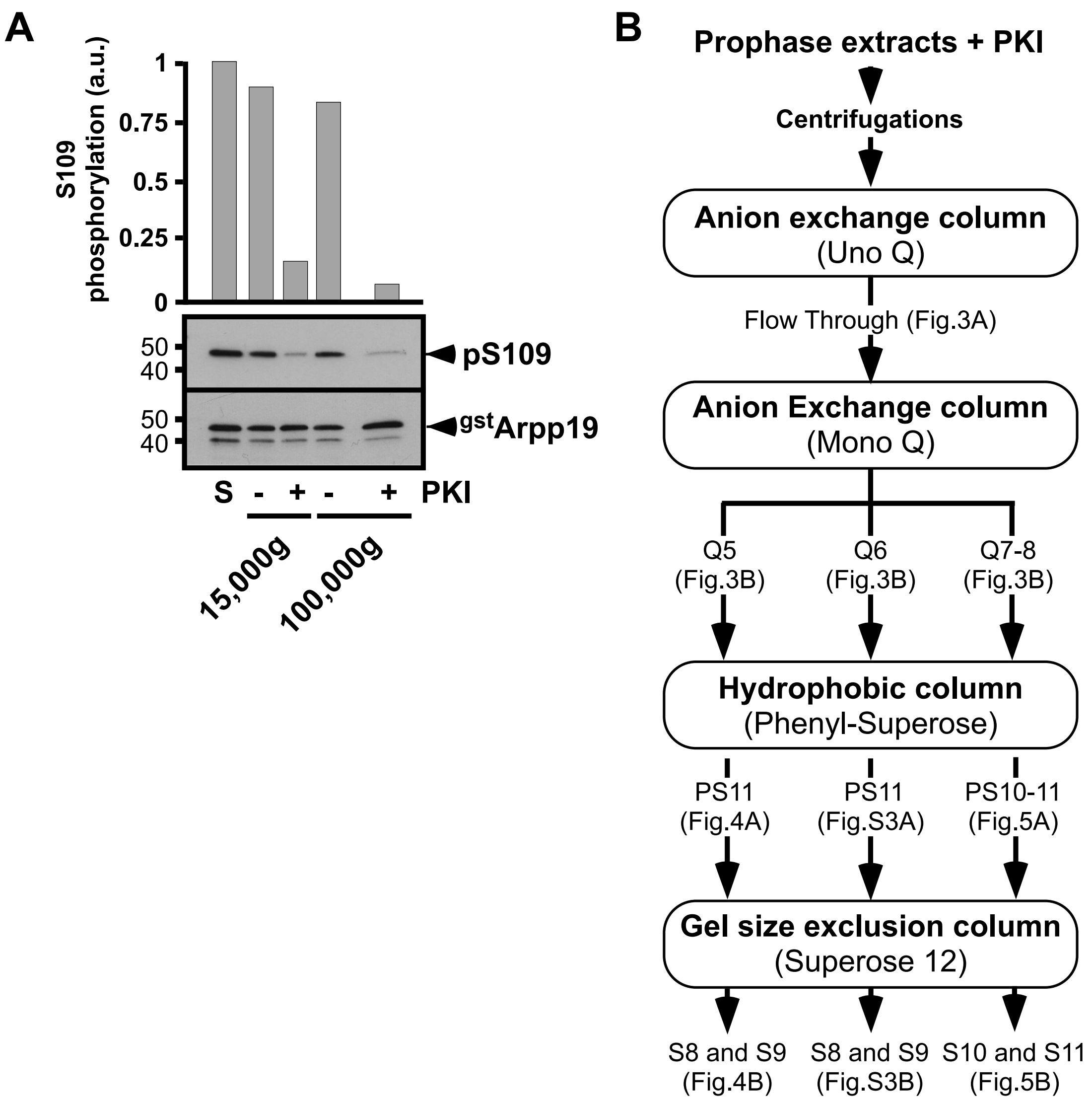


(**A**) 20,000 prophase oocytes were lysed and successively centrifuged at 300g, 1100g and 15,000g for 20 min. The 15,000g supernatant was ultracentrifuged at 100,000g for 2h. S109-phosphatase activity was assayed in the 15,000g and 100,000g supernatants by adding pS109-GST-Arpp19 in the presence or in the absence of PKI. S109 phosphorylation of GST-Arpp19 (pS109) and total GST-Arpp19 ($^{gst}$Arpp19) were western blotted using respectively phospho-S109-Arpp19 and GST antibodies. "S": starting pS109-GST-Arpp19 substrate. S109 phosphorylation quantification: an arbitrary unit (a.u.) of 1 was attributed to the phosphorylation level of S. (**B**) Scheme of the protocol used for the biochemical isolation of S109-phosphatase illustrated in Figs. 3-5 and Supplementary Fig. 3.

**Supplementary Figure 3. Biochemical isolation of S109-phosphatase from prophase extracts - Separation of fraction 6 from the Mono Q column with Phenyl-Superose and Superose 12 columns.**

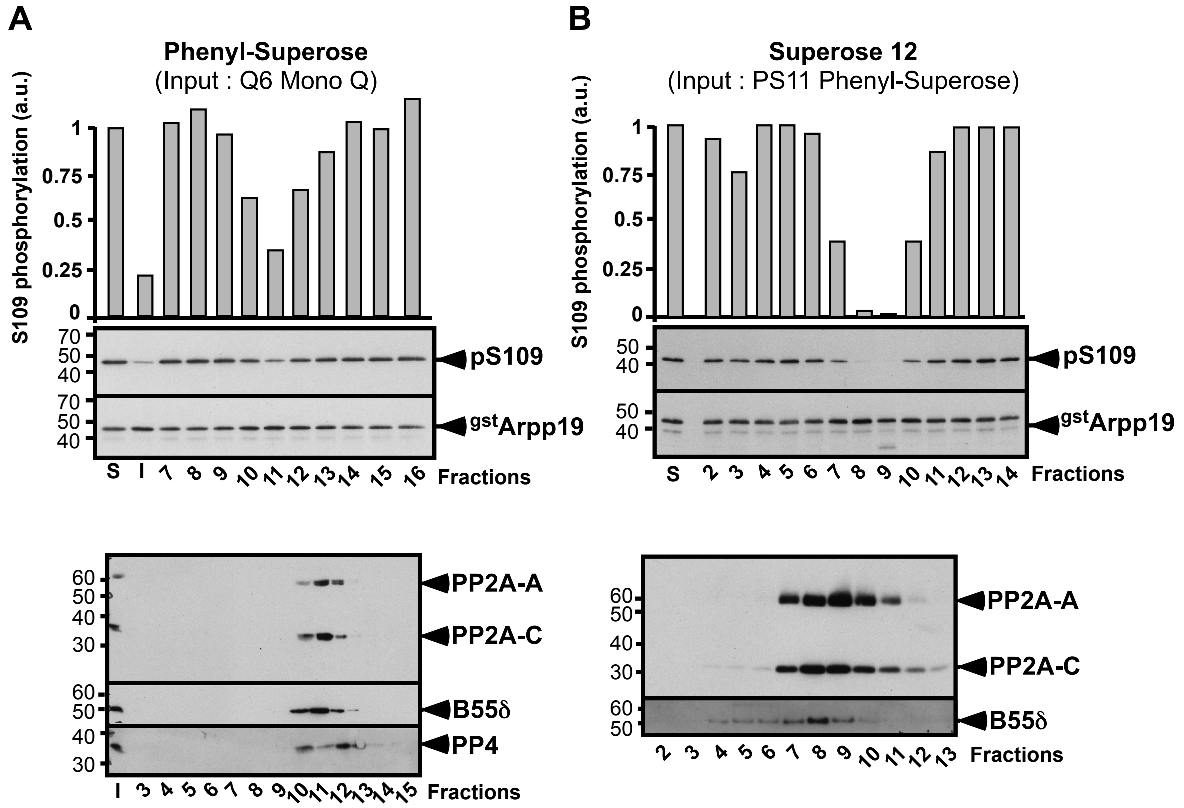


Continuation of experiment illustrated in Fig. 3. "S": starting pS109-GST-Arpp19 substrate. "I": input sample loaded on the column. S109 phosphorylation quantification: an arbitrary unit (a.u.) of 1 was attributed to the phosphorylation level of S. (**A**) Phenyl-Superose. Fraction 6 from the Mono Q column (see Fig. 3B) was loaded on the column. Elution profile of S109-phosphatase activity after Phenyl-Superose column and western blot analysis of fractions 3 to 15 with antibodies directed against catalytic subunits of PP2A (PP2A-C) and PP4, PP2A scaffold subunit A (PP2A-A) and PP2A regulatory subunit B55δ. (**B**) Superose 12. Fraction 11 from the Phenyl-Superose column (see **A**) was loaded on the column. Elution profile of S109-phosphatase activity after Superose 12 column and western blot analysis of fractions 2 to 13 with antibodies directed against PP2A scaffold subunit (PP2A-A), PP2A catalytic subunit (PP2A-C) and PP2A regulatory subunit B55δ.

**Supplementary Table 1. LC-MS/MS analysis of the S109-phosphatase enriched fractions from two distinct isolation experiments.**

The isolation procedure of S109-phosphatase was repeated 4 times with oocytes collected from different females. The results of the experiment 1 are shown in Table 1. Experiment 4 was analyzed by western blot but not by LC-MS/MS. In experiment 2, S109-phosphatase activity was recovered in fractions Q5 to Q8 after the Mono Q column. These fractions were pooled before loading on Phenyl-Superose column. In experiment 3, S109-phosphatase activity was recovered in a single fraction after the Mono Q column, Q8, which was further loaded on the Phenyl-Superose column. For both experiments 2 and 3, S109-phosphatase activity was recovered in PS11 after the Phenyl-Superose column. PS11 was loaded on the Superose 12 column. The phosphatase content estimated by LC-MS/MS sequencing in PS11 of experiment 3 and in fractions from Superose 12 column of experiments 2 and 3 is summarized in (**A**) and (**B**) for experiments 2 and 3 respectively. S109-phosphatase activity is indicated with "+" or "-". When S109-phosphatase is present, the column is in grey.

(**A**) LC-MS/MS analysis of Superose fractions from experiment 2 (input: pool of Q5 to 8 > PS11)

| | | | | | Mascot scores | | | |
|---|---|---|---|---|---|---|---|---|
| **Family** | **Description** | **Accession** | **Gene Symbol** | **Coverage (%)** | **S5** | **S7** | **S8** | **S9** |
| **PP2A-A** | Regulatory subunit A, β | 148230849 | ppp2r1b; ppp2r1b.S | 60 | 810 | 2675 | 2392 | 1509 |
| | 65 kDa regulatory subunit A β isoform-like | 148227844 | LOC398563 | 58 | 829 | 2625 | 2458 | 1509 |
| | Regulatory subunit Aα | 148222150 | ppp2r1a-b; ppp2r1a.S | 52 | 650 | 1380 | 1566 | 706 |
| | Regulatory subunit A α | 148224496 | ppp2r1a-a; ppp2r1a.L | 52 | 562 | 1205 | 1438 | 764 |
| **PP2A-C** | Catalytic subunit, α | 148230509 | ppp2ca; ppp2ca.L | 56 | 399 | 1332 | 1014 | 472 |
| | Catalytic subunit β [X. tropicalis] | 53749698 | ppp2cb | 53 | 399 | 1306 | 1003 | 472 |
| **B55** | 55 kDa regulatory subunit Bδ | 147900119 | ppp2r2d; ppp2r2d.S | 34 | 65 | 171 | 296 | 233 |
| | phosphorylase phosphatase/B55α | 963087 | ppp2r2a; ppp2r2a.S | 33 | 66 | 324 | 403 | 427 |
| **B56** | Regulatory subunit B'α | 168693595 | ppp2r5a; ppp2r5a.S | 51 | 317 | 761 | 707 | 404 |
| | Regulatory subunit B'ε | 148236023 | ppp2r5e; ppp2r5e.S | 32 | 35 | 226 | 181 | 126 |
| | | 147902694 | ppp2r5e1 | 26 | 35 | 251 | 138 | 105 |
| | Regulatory subunit B'γ | 148236119 | ppp2r5c; ppp2r5c.L | 8 | | 105 | 70 | 43 |
| **PP2C** | PP1A | 147905165 | ppm1a; ppm1a.L | 26 | | | | 184 |
| | Mg2+/Mn2+ dependent, 1B | 148227634 | ppm1b; ppm1b.L | 18 | | | | 164 |
| **PP1** | Catalytic subunit, α | 147903539 | ppp1ca; ppp1ca.L | 9 | 67 | 35 | 34 | 36 |
| **PP3** | Catalytic subunit, α | 148235616 | ppp3ca | 9 | 82 | | 62 | 108 |
| **PP4** | Catalytic subunit [Xenopus tropicalis] | 45360541 | ppp4c | 13 | | 113 | 60 | 37 |
| **PP6** | Catalytic subunit | 147906292 | ppp6c; ppp6c.L | 3 | 23 | | | |
| **S109-phosphatase activity** | | | | | **-** | **-** | **+** | **-** |

(**B**) LC-MS/MS analysis of PS11 and Superose fractions from experiment 3 (input: Q8 > PS11)

| | | | | | **Mascot scores** | | | | | | |
|---|---|---|---|---|---|---|---|---|---|---|---|
| | | | | | | **Superose fractions** | | | | | |
| **Family** | **Description** | **Accession** | **Gene symbol** | **Coverage (%)** | **PS11** | **S5** | **S6** | **S7** | **S8** | **S9** | **S10** |
| **PP2A-A** | 65 kDa regulatory subunit Aβ isoform-like | 148227844 | LOC398563 | 52,63157895 | 1888 | 388 | 251 | 415 | 620 | 838 | 403 |
| | Regulatory subunit Aβ | 148230849 | ppp2r1b; ppp2r1b.S | 52,12224109 | 1825 | 347 | 255 | 395 | 572 | 814 | 403 |
| | Regulatory subunit Aα | 148222150 | ppp2r1a-b; ppp2r1a.S | 42,44482173 | 1181 | 141 | 51 | 290 | 370 | 454 | 218 |
| | Regulatory subunit Aα | 148224496 | ppp2r1a-a; ppp2r1a.L | 41,93548387 | 1135 | 141 | 51 | 321 | 399 | 484 | 218 |
| **PP2A-C** | Catalytic subunit α | 148230509 | ppp2ca; ppp2ca.L | 53,07443366 | 721 | 171 | 69 | 108 | 214 | 227 | 99 |
| | Catalytic subunit β [X. tropicalis] | 53749698 | ppp2cb | 50,48543689 | 690 | 155 | 69 | 132 | 233 | 243 | 99 |
| **B55** | Regulatory subunit Bα | 148234757 | ppp2r2a; ppp2r2a.S | 46,17117117 | 866 | 56 | 67 | 106 | 81 | 151 | 42 |
| | 55 kDa regulatory subunit Bδ | 147900119 | ppp2r2d; ppp2r2d.S | 27,74049217 | 706 | 65 | 88 | 88 | 66 | 71 | 33 |
| **B56** | Regulatory subunit B'α | 168693595 | ppp2r5a; ppp2r5a.S | 23,52941176 | 438 | | | | | 43 | |
| | Regulatory subunit B'β | 148234627 | ppp2r5b; ppp2r5b.L | 26,2605042 | 452 | 41 | | | | 43 | |
| | Regulatory subunit B'ε | 148236023 | ppp2r5e; ppp2r5e.S | 18,84368308 | 268 | | | | | | |
| | | 147902694 | ppp2r5e1 | 17,55888651 | 249 | | | | | | |
| | Regulatory subunit B'γ | 148236119 | ppp2r5c; ppp2r5c.L | 4,347826087 | 30 | | | | | | |
| **PP2C** | PP1A | 147905165 | ppm1a; ppm1a.L | 46,73629243 | 1011 | | 41 | 31 | | 218 | 389 |
| | Mg2+/Mn2+ dependent, 1B | 148227634 | ppm1b; ppm1b.L | 15,42168675 | 549 | | 24 | | | 81 | 139 |
| **PP1** | Catalytic subunit Bγ | 148224548 | ppp1cc | 23,043 | 206 | | | | | | |
| **PP3** | Catalytic subunit, α | 148235616 | ppp3ca | 5,598455598 | 94 | | | | | | |
| **PP6** | Catalytic subunit | 147906292 | ppp6c; ppp6c.L | 10,16393443 | 15 | | | | | | |
| **PP4** | Regulatory subunit 2-B | 147901735 | ppp4r2; ppp4r2.S | 3,768844221 | 19 | | | | | | |
| **S109-phosphatase activity** | | | | | +++ | - | + | ++ | ++ | + | - |
